# Advancing source tracking: systematic review and source-specific genome database curation of fecally shed prokaryotes

**DOI:** 10.1101/2024.06.14.598882

**Authors:** Blake G. Lindner, Rakin A. Choudhury, Princess Pinamang, Lilia Bingham, Isabelle D’Amico, Janet K. Hatt, Konstantinos T. Konstantinidis, Katherine E. Graham

## Abstract

Approaches for shotgun metagenomic sequencing are being adapted to many environmental monitoring tasks but remain poorly suited for source attribution within fecal source tracking (FST) studies due to a lack of knowledge regarding the source specificity of fecally shed microbial populations. To address this gap, we performed a systematic literature review and curated a large collection of genomes (n=26,018) representing fecally shed prokaryotic species across broad and narrow source categories commonly implicated in FST studies of recreational waters (i.e., cats, dogs, cows, seagulls, chickens, pigs, birds, ruminants, human feces, and wastewater). These data include genome sequences recovered from metagenomic and isolation-based studies which we examined extensively with comparative genomic approaches to characterize trends across source categories and produce a finalized genome database for each source category which is available online (n=12,730). We find that across these sources, the total number of genomes recovered varies substantially: from none in seagulls to 9,085 in pigs.

According to available data, on average 81% of the genomes representing species-level populations occur only within a single source. Using fecal slurries to test the performance of each source database, we report read capture rates that vary with fecal source alpha diversity and database size. Lastly, while extensive work has been performed on the rumen system of cows and other ruminants, we note the scarcity of genomic characterization of prokaryotes in the feces of these animals. We expect this resource to be useful to FST-related objectives, One Health research, and sanitation efforts globally.

**Table of Contents Graphical Abstract:** 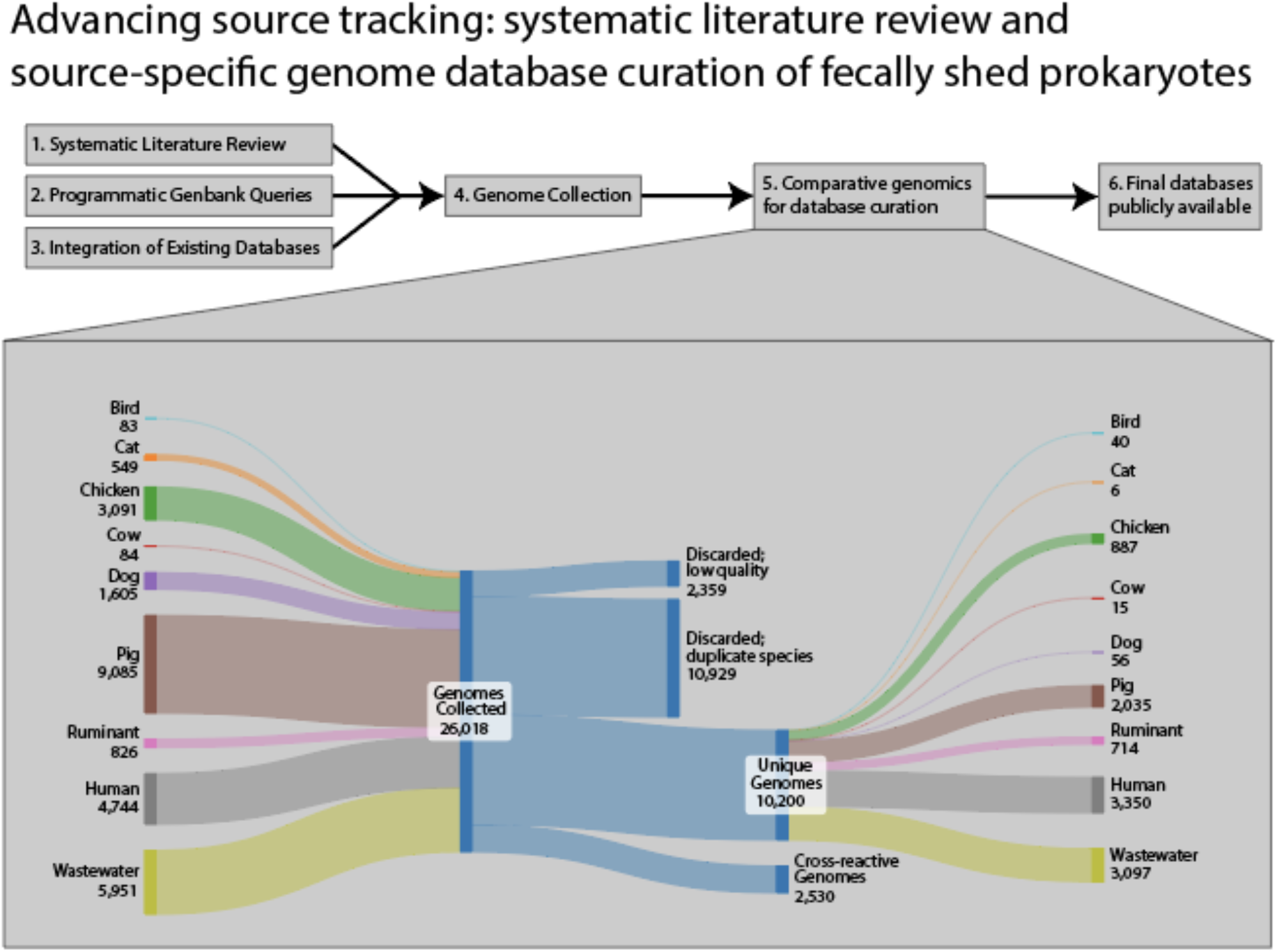

**Synopsis:** We curate nine prokaryotic genomic databases for fecal source-tracking, summarize their source-specificity, and show their usefulness by bioinformatically testing mock impaired surface water samples.

## Introduction

Globally, more than 120 million yearly cases of gastrointestinal disease result from exposure to sewage-impacted surface waters used for swimming and bathing.^1^. Health risks due to sewage contamination in the environment are assessed using fecal indicator bacteria (FIB).^2^ However, studies have shown that FIB can be insufficient for assessing fecal pollution in a given waterbody, as they are non-conservative and non-specific.^3–5^ These issues have borne the field of microbial source tracking (MST) or fecal source tracking (FST), which seeks to identify sources of fecal pollution in the environment, evaluate health risks, and guide remediation.^6,7^

Historically, FST methods have focused on using PCR- or qPCR-based approaches to quantify source-specific genetic markers in samples from impaired waters to infer the presence of sewage. Genetic markers within the 16S rRNA gene of *Phoecaeicola dorei* (formerly *Bacteroides dorei*), such as HF183,^8,9^ or viral genetic markers, such as for crAssphage have been used with success to identify sewage in the environment.^10^ However, this approach can suffer from a lack of source-specificity.^11,12^ 16S rRNA gene sequencing has emerged as a potentially useful tool in the MST toolbox, with mixed results to date.^13,14^ A more robust characterization of source-specific populations may assist with identifying genetic signatures that can more robustly attribute fecal sources in impaired waters – representing another approach in the FST methodological toolbox.

The advent of metagenomics has allowed for the characterization of prokaryotic genomes across many different microbiomes, especially the human gastrointestinal system. Datasets representing comprehensive efforts to collect species-level genomic representations of the prokaryotic organisms inhabiting gastrointestinal microbiomes are growing in size and frequency,^15–17^ covering an increasing number of animals. Yet, no database exists which collates this emerging data into a single repository for use within an FST context. Leveraging publicly available data from isolation-based and metagenomic studies can reveal genomes and gene content shared between sources of fecal contamination thus helping identify source-specific features that could drive future innovation among FST methodologies^18^.

To this end, this systematic review and meta-analysis aimed to gather previously published datasets from metagenomic or isolate studies associated with prokaryotic extraintestinal fecal shedding from the following animal sources for database curation: cow, pig, chicken, cat, dog, goat, seagulls, birds (non-seagull), and ruminants (non-cow). We then assessed ecological questions regarding the genomes: What genomes are specific to each source? How well do collections of genomes capture fecal signal? In parallel, we collected, and shotgun sequenced fecal samples from the above sources in Georgia, USA, to test the read mapping performance of our databases. These efforts can be informative to practitioners of MST and those interested in developing new MST genetic marker assays based on fecally shed prokaryotic populations.

## Materials and Methods

### Systematic literature review, genome accessioning, and collection

A systematic literature review was undertaken to collect prokaryotic genomes from existing studies on the fecal microbiomes of the eight animal source categories. PRISMA guidelines were followed during the systematic review.^19^ Exact search strings and detailed review protocols are documented in supporting information.

Additionally, we sought to accession genomes into our collection by programmatically searching Genbank– a central genome repository managed by NCBI/DDBJ/ENA – independent of searching published literature. Genbank search details and collected genomes are provided in the SI. We also collected large genome databases stored elsewhere (e.g., Zenodo, EMBL, CNDB) from noteworthy publications or major studies produced after or missed by our initial literature review (particularly for human and wastewater sources.^20–23^ Further information on these methods can be found in supporting information.

All genomes gathered through the above methods are recorded in the supporting information with relevant metadata and download sources for reproducibility (Tables S2 and S3).

### Comparative genomics and database curation

We constructed a software pipeline to process the genomes collected through our efforts. Documentation, scripts, and installation instructions are hosted online (https://github.com/blindner6/SourceApp). Additionally, a detailed summary of our workflow can be found in the supporting information. In brief, we used comparative genomic approaches to identify and discard genomes of poor quality, organize remaining genomes into species-level clusters (≥95% ANI), and flag genomes belonging to different sources but assigned to the same species-level cluster as cross-reactive.

Our database is available for download online (https://doi.org/10.5281/zenodo.10728776).

### Fecal slurry creation, DNA extraction and Illumina Sequencing

To characterize the performance of our genome databases, we collected fecal samples from animals that may contribute to surface water contamination in Atlanta, GA, USA. Fecal sources in this study included: cow and pig (from one farm), dog and cat (from one shelter), as well as chicken and goat (from one farm). We also collected 1 L of 24-hour-composited primary influent samples from three water reclamation facilities, and 500 mL of composited septage from a septage pumping truck. We created fecal slurries by homogenizing fecal samples from ten individuals of each source (dog, cat, chicken, goat, pig, cow) from which we extracted DNA and shotgun sequenced approximately 4.5×10^9^ bp (Gbp) for each. With equal amounts of sequencing effort (∼4.5Gbp) applied to each slurry, we assessed the fraction of sequence diversity (an approximation of alpha diversity) observed by the given sequencing effort via Nonpareil.^24^ Controlling our experiments in this way resulted in estimations for the coverage of a slurry’s sequence diversity that varied based only on community complexity (i.e., sequence diversity).

Additional details on the creation, sequencing, and analysis of these samples can be found in the supporting information. Short reads associated with the fecal slurries were deposited to NCBI under BioProject PRJNA1092107.

## Results and Discussion

### Systematic literature review and data collection

The systematic literature review revealed trends in efforts across the world to characterize animal fecal prokaryotes at the whole-genome level. Several sources had few numbers of studies producing prokaryotic genomes from feces, such as seagulls (0 studies) and cats (n=1 study.

Birds (ducks, starlings, geese, finches, starlings, wigeons, swallows, crows, pheasants), cows, pigs, and ruminants, each had more than 10 studies (Table S1). The systematic literature review also revealed many different sequencing and bioinformatics methods used for data processing and genome binning. Culture-based methods were the most common in literature. Some studies used shotgun metagenomes (a culture independent approach) to recover genomes, but this approach was far less common. Most studies used short-read platforms (Illumina) to generate sequencing data, but some used long-read platforms (PacBio or ONT) either individually or as a hybrid sequencing approach.

### Curation and comparative genomics of source databases

We collected 26,018 genome sequences representing fecally shed prokaryotic populations across the selected sources. Figure 1 summarizes the number of genomes collected for each source, their outcomes following our examination, and to what extent genomes were unique to a source. Following our curation efforts, 2,359 genomes were discarded due to poor quality scores. An additional 10,929 genomes were discarded during species-level clustering (≥95% ANI^25^), which aims to leave only a single representative genome for each species-level cluster. Thus, the remaining 12,730 genomes represent species found in each of our sources. Of these genomes, 2,530 (19%) represent species observed in multiple sources (i.e., cross-reactive species; Figure 1C).

**Figure 1.**
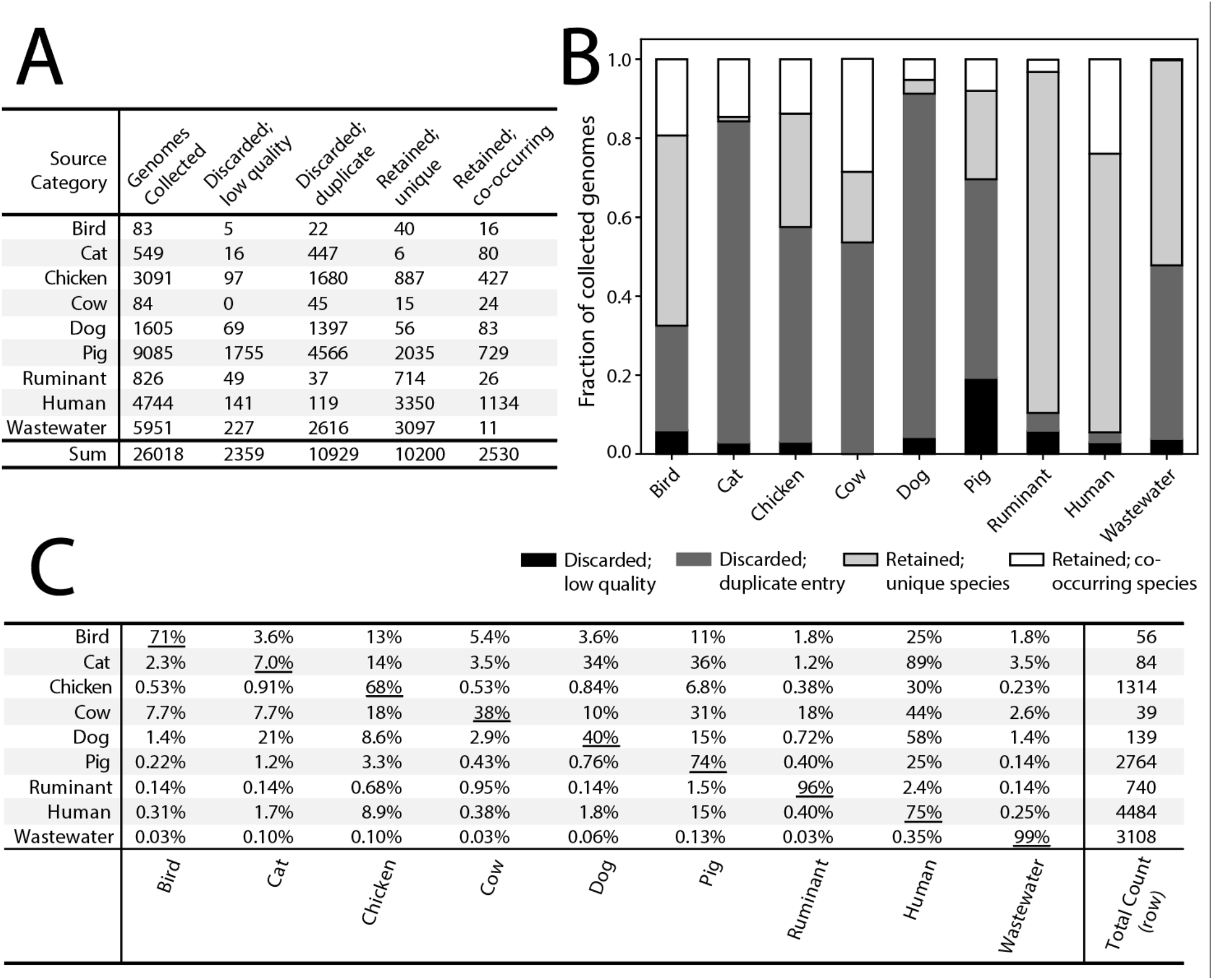
Overview of genome search results and curation of source databases. (A and B) The total genomes found for each source and their outcome in the study. (C) Matrix summary of the fraction of genomes representing shared species across source categories. Underlined values on the diagonal denote the fraction of source-specific genomes for a given row.

The fraction of species in the curated database specific to a given source varied substantially across sources. For example, the human genome collection contained the highest number of cross-reactive species (25% or 1,134) but was also the collection with the greatest number of species represented altogether (n=4,484 species). The much smaller cat and dog collections exhibited more cross-reactivity with 93% of cat species (n=77) and 60% of dog species (n=81) co-occurring in other sources. In this instance, nearly all the cross-reactive cat and dog species were also found in the human database. In contrast, the wastewater collection exhibited virtually no cross-reactivity with any other source (11 species; 0.4%). In general, the largest numbers of shared species were between either (1) the bird, cat, chicken, cow, dog, and pig databases with the human data, or (2) between source categories that are closely related in habitats (e.g., dogs and cats).

We summarized the location metadata associated with all genomes collected regardless of their inclusion in the finalized source databases to show the geographic breadth of the data collected through our efforts (Figure 2A). Most genomes originated from China (n=9780) and the USA (n=4274), though many genomes (n=3674) had no metadata describing their geographic origin. It is crucial to note that our efforts returned comparatively few genomes from countries in the Southern Hemisphere.

**Figure 2.**
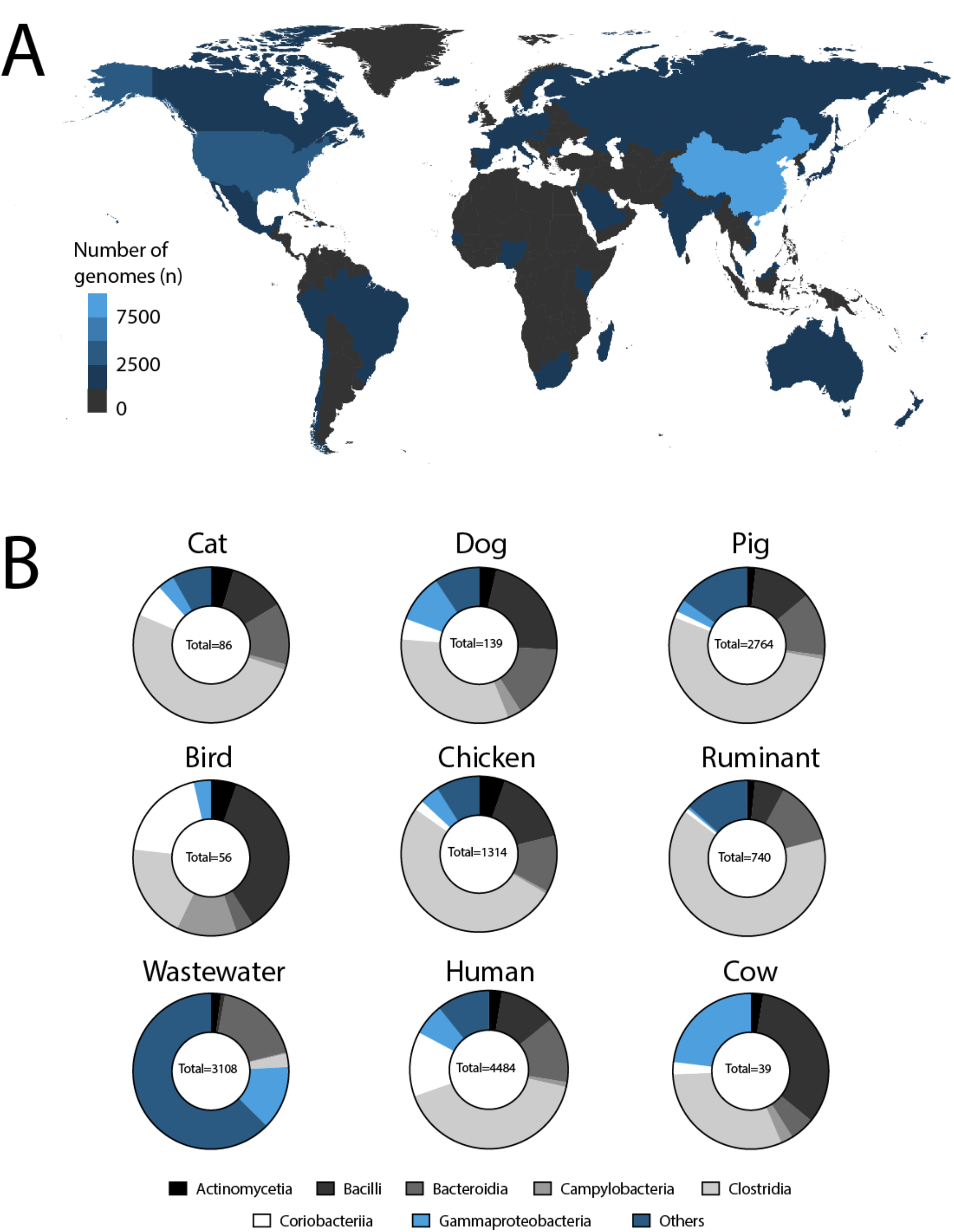
(A) Geographic distribution of genomes (quality scores ≥ 50) and (B) number of species within a taxonomic Class accessioned within the MST genome database (n=12,730 genomes). The top seven Classes are shown while remaining Classes are grouped together under “Others”.

We examined common and unique taxonomies across source categories, as we dereplicated genomes within source databases at the species level (Table S4). Gut-associated classes, such as *Actinomycetia, Gammaproteobacteria, Bacilli, Clostridia, Campylobacteria, and Bacteroidia*, were common across all source categories. The larger genome databases also had unique taxonomies (e.g., human, wastewater), likely owing to their more complete characterization relative to the underexplored sources (e.g., cow). For instance, *Desulfitobacteriia, Halobacteria*, and *Thermoanaerobacteria*, were uniquely found in the human fecal genome database. The wastewater database had 157 unique Classes compared to the other source category databases.

### Assessing database performance

Laboratory assembled fecal slurries, primary influent, and septage were filtered, extracted, sequenced, and the diversities of their resulting metagenomes compared across sources. Values for the percentage of observed sequence diversity (or Nonpareil coverage^24^) ranged from 46% (cow) to 97% (cat) (Figure 3, Top; Table S5). These results reveal that for more complex slurries (e.g., cow, goat), substantially greater amounts of sequencing effort are required to cover most of the community’s diversity. For example, approximately 1-4 Gbp of sequencing effort would be required to observe 95% of the sequence diversity for the least diverse fecal slurries (cat and dog) whereas approximately 400 Gbp would be required to achieve that level for the most diverse fecal slurry (cow). These results can inform the planning of studies utilizing assembly and binning of shotgun metagenomic data to recover new genomes for a particular source category^26^.

**Figure 3.**
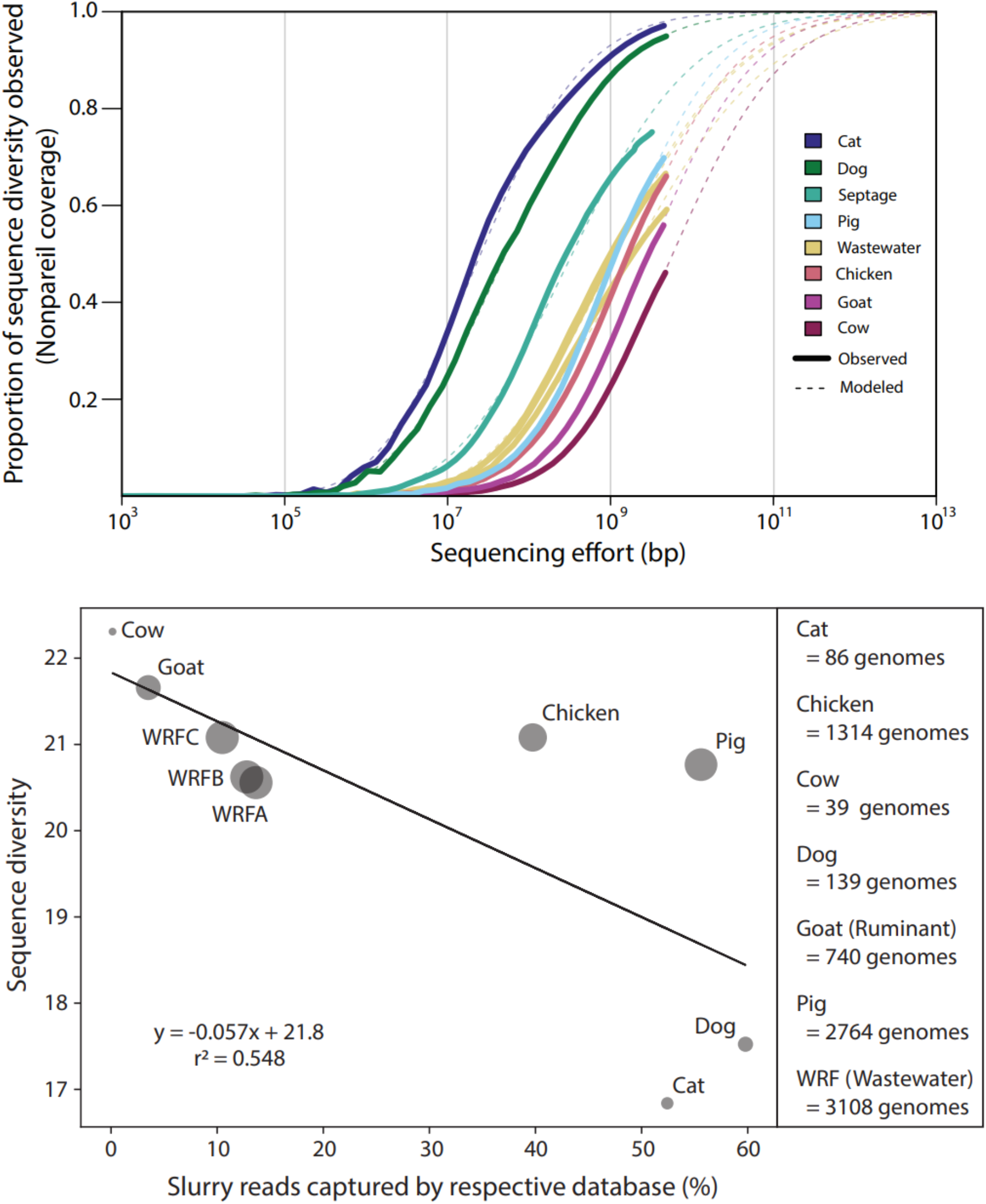
Fecal slurry sequencing results and genome database performance. (Top) The fraction of sequence diversity observed (y-axis) for each fecal slurry as estimated by Nonpareil shown as functions of sequencing effort (x-axis). The underlying data provided to Nonpareil were short read metagenomes of fecal slurries from source categories used in this study. Solid lines represent the observed relationship and dotted lines represent extrapolated models; approximately 4.5 Gbp of sequencing effort per sample was used. (Bottom) Nonpareil diversity for each slurry as a function of reads mapped from the slurry to its respective curated genome database. The size of each point is scaled to represent the number of species-level dereplicated genomes represented in the respective source. The number of species-level dereplicated genomes present in each source database are shown for reference in the bottom right-hand panel.

To understand the capabilities of each source’s curated database in an FST context, we mapped the resulting short read metagenomes to their respective genome databases (i.e., dog fecal slurry short reads queried against the dog database). In this way, we aimed to assess what fraction of the underlying short-read data our curated databases could capture, with the expectation that higher rates of read capture imply a more representative database. When plotting sequence diversity as a function of slurry read capture by their respective databases, a linear relationship was found with a negative slope (Figure 3, bottom). This relationship implies that read capture rates are inversely proportional with sequence diversity. The dog and cat databases underperformed expectations based on this relationship and the chicken and pig databases overperformed, though both sets of databases vary substantially in terms of size. Thus, an additional variable may be crucial for understanding these trends i.e., the number of species represented within each source database. Both chicken and pig databases are relatively speciose with most species appearing source-specific (Figure 2) while dog and cat contain drastically fewer species. Consequently, we hypothesize that – independent of fecal sequence diversity – as a source’s genome database becomes increasingly comprehensive, read capture rates will increase concomitantly.

Next, to understand the degree to which these source databases manage instances of cross-reactivity, we competitively mapped each fecal slurry metagenome against every source database to obtain new read capture rates (Figure 4; Table S6). These results did not include assembly and binning of the fecal slurry metagenomes themselves, as the goal was to test the efficacy of each source database without the addition of de novo references (i.e., new MAGs) from the slurries. For most fecal slurry metagenomes, its respective genome database was the dominant database against which reads were mapped except for the cow and goat slurries. We expected the goat slurry to map predominantly to the ruminant database, however, it best mapped to the chicken database. Some cross-reactivity from fecal metagenomes to non-matching genome databases was observed, especially for cat and dog metagenomes, as they frequently mapped to each other and to the human database. The effect of shared facilities for some of the sources may have contributed to some of the overlap we observed in mapped reads among these sources (e.g., dog and cat, chicken and goat). This limitation should be addressed in future efforts.

**Figure 4.**
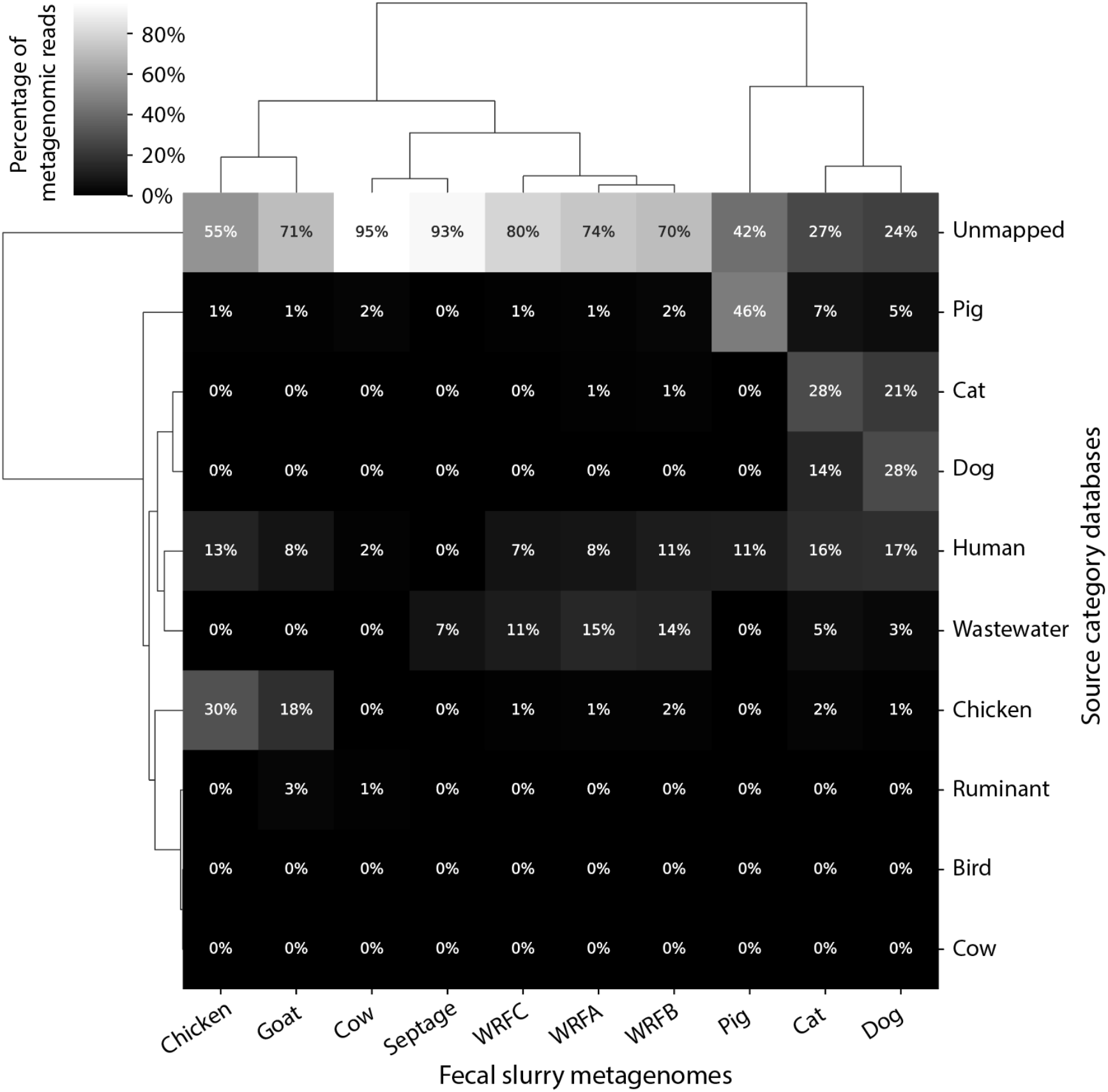
Competitive read mapping performance of metagenomic reads from experimental slurries (columns) prepared in the laboratory mapped to our curated genome databases (rows). Both rows and columns have been clustered to minimize variance within groups using Ward’s method for the values shown. Reads were competitively mapped to genomes within databases and summed within each database to obtain the percentage of reads mapped to each database. WRF = wastewater reclamation facility.

### Relevance for environmental monitoring

Ongoing efforts to catalog prokaryotic diversity across fecal sources are occurring rapidly but have yet to be synthesized into a single database with proven usefulness for FST. This study provides the most comprehensive genomic databases to date cataloging fecally shed prokaryotes from commonly implicated non-human sources responsible for surface water impairment. This work explored source specific database construction as the necessary next step for metagenomic sequencing and its associated bioinformatic approaches to inform FST in a practical manner. We have automated the procedure for constructing these databases which will facilitate future expansion as more genome sequences become available. Lastly, our fecal slurry metagenome read mapping exercises against these source databases represent a relatively simple experimental approach to assess the efficacy of metagenomic approaches for FST *in vitro* and inform potential users of its possible strengths and weaknesses.

Despite the predominance of genomes produced from culture-based workflows (e.g., isolation/enrichment studies) in our datasets, our results are broadly consistent with 16S rRNA gene sequencing studies documented in the literature for FST. For instance, Li et al. used a microarray platform and 16S rRNA gene amplicon sequencing to characterize both sewage and avian, cattle, poultry, swine feces and found that families *Ruminococcaceae* and *Lachnospiraceae* were common amongst all groups.^27^ Our results were consistent with these findings and we identified more classes that were common across these sources and in new sources including dogs and cats (e.g., *Actinomycetia*). Partial 16S rRNA gene sequences are not, however, robust species-level delineators,^28^ though others have reported the use of 16S rRNA gene sequencing alone for FST efforts to attribute fecal pollution to specific sources.^14,29^ Several bioinformatic tools exist that utilize sequencing data for such analyses, including FEAST,^30^ FORENSIC,^31^ and (meta)SourceTracker for which our database may be useful.^32,33^ Each of these tools uses underlying probabilistic or statistical models to model the inputs of sources to sinks.

As such, sources must be provided with sink datasets to characterize the attributions of sources to each sink sample, which are not always readily available when working on issues of surface water contamination from point or non-point sources.

The human gut microbiome represents the best studied fecal source with work summarizing our understanding of its extant species diversity estimating it is probably comprised of about 3,000 reoccurring (i.e., excluding singletons) species globally^20^. Work examining human feces at the metagenomic level previously reported sequence diversity values less than those we found for our wastewater samples but greater than those of the dog and cat fecal slurries^34,35^.

Unsurprisingly, this suggests there are a substantial number of species missing from the databases of many of the sources we examined. Indeed, no source saw read capture rates exceed 60% in our read mapping exercises (Figure 3). Considering the risk implications of cow-derived fecal contamination in recreational waters and agriculture^36^ and the relatively few publicly available genomes from cow feces (n=84), this work highlights the need for more substantial research efforts to understand microbial communities in cow feces, as opposed to the rumen alone (Figure 1). Our work highlights the need for more global surveys of fecally shed prokaryotes across major sources to address issues of variation in fecal diversity among individuals of the same source (e.g., biodiversity; Figure 2). Issues of internal variation may complicate interpreting the results of shotgun metagenomic results in FST work and should be addressed in future work.

As we have seen above, this approach will suffer when a source has a high degree of sequence diversity but is poorly characterized (Figure 3). For example, we observed the largest percentage of fecal slurry metagenomic reads typically mapped to their same database in the competitive mapping exercises except in the case of cows and ruminants (Figure 4). In terms of other limitations, our databases utilized only assembled genomes, which greatly limited the publications that met our inclusion criteria. We recommend that authors using pipelines that result in assembled sequences upload both their assemblies and raw reads to facilitate the curation of databases such as this in the future. In the future, database sizes can be increased for each source by downloading and assembling publicly available metagenomes and before augmenting our databases. Finally, in their current state, the source-specific databases are far from complete, owing to the relatively limited information we have on animal fecal microbiomes; therefore, we caution against drawing broad biological or ecological conclusions from our summaries above. For instance, there were no publicly available assemblies from the feces of seagulls, although they are often a source of pollution to surface waters and can carry human pathogens.^37–39^. This work highlights the need for ongoing research on animal fecal microbiomes across many species, geography, and lifestyles, as well as septage to inform microbial source tracking efforts and One Health research.

## Supporting Information

Supporting information for this work includes:

- Additional methods for systematic literature review, database construction, and comparative genomics.
- Summaries of systematic literature review results, genome collection, and results of comparative genomic analyses.

## Acknowledgments

This work has been supported by the US National Science Foundation (Award No 2136146), the US EPA (Award No 84020301-0), and the Georgia Institute of Technology’s President’s Postdoctoral Fellowship Program.

