## Supplemental Methods for "Advancing source tracking: systematic review and source-specific genome database curation of fecally shed prokaryotes"

### Additional Materials and Methods

#### Systematic literature review

The goal of the review was to first gather prokaryotic genomes representing populations within animal stool which are shed into the environment, then bioinformatically eliminate cross-source genomes, and finally produce source-specific genomic fecal indicators. The review inclusion criteria were: (1) the article must be written in English, (2) the article must not be a review and must present peer-reviewed primary data, (3) the article describes methods that are justifiable, (4) the sampling approach must focus on fecal or stool samples (i.e., rather than rumen, swabs), (5) the article must provide metagenome-assembled genomes (MAGs) or isolate genomes (rather than metagenomes alone; no assembly and binning of any metagenome took place as part of this study), (6) the sampling and laboratory processing approach must yield prokaryotic genome sequences (as opposed to exclusively viral or eukaryotic), (7) the hosts must be wild or domesticated animals (as opposed to laboratory animals fed experimental diets and/or are specific-pathogen free), and (8) due to their prevalence in the literature and cross-species presence, we made efforts to exclude genomes that were zoonotic pathogens (*Salmonella enterica, Listeria monocytogenes, Escherichia coli, Campylobacter jejuni, Klebsiella pneumoniae, Yersinia pestis, Clostridium perfringens, Enterococcus faecalis, Mycobacterium avium, Clostridium difficile, Enterococcus faecium, Campylobacter coli, Salmonella typhi*). We did not focus on any human sources in our review, as there are already robust efforts to summarize prokaryotes of the human gut.[1,2]

Table 1: Search strings used for each source category. Strings were queried on the Web of Science and Scopus databases on 27 January 2023, and the results were exported and input into Rayyan for title review and abstract review phases.

| Source category | Search string |
|---|---|
| *Cow* | (cow OR bovine OR bos OR bov*) AND (mags OR genomes) AND (fec* OR stool OR faecal) |
| *Pig* | (pig OR sus OR porcine OR swine) AND (mags OR genomes) AND ABS (fec* OR stool OR faecal) |
| *Chicken* | (chicken OR gallus OR poultry) AND (mags OR genomes) AND (fec* OR stool OR faecal) |
| *Cat* | (cat OR feline OR felis) AND (mags OR genomes) AND (fec* OR stool OR faecal) |
| *Dog* | (dog OR canis OR canid OR canine) AND (mags OR genomes) AND (fec* OR stool OR faecal) |
| *Seagull* | (seagull OR gull) AND (mags OR genomes) AND (fec* OR stool OR faecal) |

| | |
|---|---|
| *Ruminant (non-cow)* | (ruminant OR goat OR deer OR sheep OR antelope OR giraffe OR gazelle) AND (mags OR genomes) AND (fec* OR stool OR faecal |
| *Bird (non-gull)* | (bird OR avian OR aves) AND (mags OR genomes) AND (fec* OR stool OR faecal) |

Search strings for each source category were queried against the Web of Science and Scopus databases, and records were imported to Rayyan for review. Records were deduplicated and title review proceeded, followed by abstract review, then full text review, and finally genome extraction. During each review stage (title, abstract, and full text), two reviewers were assigned to each source, and each reviewer was blinded to the other's decision regarding inclusion/exclusion from the review. Once both reviewers were finished with a stage, their decisions were unblinded and for the publications that had conflicting inclusion/exclusion decisions between reviewers, the reviewers met to discuss a consensus decision before proceeding to the next phase. We gathered accession numbers for genomes recovered from publications that passed the full text review stage of our systematic literature review. This was accomplished via accessioning numbers (in NCBI, Zenodo) provided in the publication or correspondence with the authors of the study. In either of these instances, we further checked all available metadata for genomes to ensure they still met our inclusion criteria. This effort was useful in instances where the exact information about the genome was clearer from database metadata than publication text. In very rare instances, metadata regarding a genome disagreed with our conclusion based on full text review. In these instances, we conservatively discarded those genomes.

Systematic genomic database search

Additionally, we sought to capture genomes that were deposited in public genomic databases but otherwise evaded our systematic literature review. Thus, the same search strings were used to programmatically query NCBI/DDBJ/ENA's Genbank databases for such assemblies via the ncbi-datasets tool (https://github.com/ncbi/datasets). This process involved the application of these search strings serially by repeated calls to "datasets summary genome taxon bacteria --search" (and similarly for "archaea") followed by each given search string. The resulting metadata describing each genome returned by our search was concatenated into a single JSON file for each source category. We then manually examined each entry according to the same criteria applied in the systematic literature review. Genomic data that met our inclusion criteria were subsequently downloaded and metadata retained for further analysis.

Lastly, publications produced after our initial literature review and/or genome collections stored across other repositories (e.g., EMBL, CNDB) were also added to each database. These ad hoc inclusions were screened following the same logic as used above: i.e., associated metadata was carefully examined to ensure that each genome met the same inclusion criteria applied elsewhere to ensure uniform results. Further attempts were made, for instance by contacting primary authors, to confirm geography and that sampling methods met the inclusion criteria. Human fecal genomes and wastewater derived genomes, which have been extensively characterized, were obtained from existing literature.[2–5]

**Comparative genomics**

To summarize our procedure in brief, all collected genomes were run through CheckM2 (v1.0.2)[6] to flag genomes of poor quality. We then ensured that all genomes met or exceeded a single copy gene (SCG) aggregate quality score of 50. This score was calculated according to Quality (Q) = SCG completeness – 5 x SCG redundancy where "completeness" and "redundancy" were measured by CheckM2. Passing genomes (Q>= 50) were then clustered and dereplicated at the species level (~95% ANI) within each source category using dRep (v3.4.2)[7] with the following parameters: "--S_ani 0.95" and "--S_algorithm fastANI". This method of dereplication results in a single genome being selected as the best representative for a species cluster. Additionally, genome quality information was forwarded to dRep from the CheckM2 results described above to facilitate the selection of the best representative genome within a species cluster. Lastly, our pipeline used dRep to flag genomes that appeared to represent the same species present across source categories (i.e. cross-reactive entries). All genomes were retained regardless of any noted cross-reactivity between source categories.

Outside of the pipeline, all resulting representative genomes for each species cluster were annotated with rgi (v6.0.3; CARD v3.2.8)[8] and then taxonomically classified against GTDB (R214) using GTDB-tk (v2.3.2).[9,10]

**Fecal sampling, slurry creation, sequencing, and analysis**

Fecal sources in this study included: cow and pig (from one farm in Georgia, US), dog and cat (from one shelter), as well as chicken and goat (from one farm). We also collected 1 L of 24 hour-composited sewage samples from three water reclamation facilities in Georgia, as well as 500 mL of composite septage samples, in autoclaved, acid-washed plastic Nalgene bottles. For the animal fecal samples, we collected at least 10 individual fecal samples from each animal source from veterinary clinics and agricultural facilities. All samples were stored on ice in the dark during transport to the lab and were processed within 24 hours of collection.

In the lab, the 10 individual fecal samples (approximately 1 g per individual) for each source category were combined to make a fecal slurry for each in a 50 mL conical and filled to a volume of 50 mL using sterile-filtered 1X PBS. Fecal slurries were diluted 1:3 in Zymo DNA/RNA Shield (cat no. R1200-25) and stored at -80°C until nucleic acid extraction. Each sample was extracted using the Qiagen PowerSoil Pro kit (cat. no. 47014) and extracts were stored at –80°C prior to Illumina sequencing.

DNA sequencing libraries were prepared from 50 ng of input DNA for each sample using the Illumina DNA prep kit with unique dual indexing as recommended by the manufacturer. Library concentrations were determined using the Qubit 1X dsDNA High Sensitivity kit (ThermoFisher Scientific) and Qubit 2.0 fluorometer and average insert size was determined by sizing using the Agilent 2100 Bioanalyzer and a High Sensitivity DNA Analysis kit. Libraries were pooled in an equimolar mixture and sequenced by the Georgia Institute of Technology Molecular Evolution Core on an Illumina NovaSeq 6000 instrument for 2 × 150-bp paired-end reads. Adapter trimming and demultiplexing were carried out on the instrument.

Finally, to examine the amount of fecal signal our efforts could recover, the resulting short-read datasets were competitively queried against the entire genome database. We indexed the final version of our genome database (n=12,730 genomes) and queried short reads against it using BWA-mem2 (2.2.1).[11] The resulting mappings were processed using coverM (https://github.com/wwood/CoverM) to estimate the 80% truncated average sequence depth (TAD80) of each genome in the database. This information was summed for each source category and the results were normalized to genome equivalents to control for differences in each community's average genome size.[12]

(4) Singleton, C. M.; Petriglieri, F.; Kristensen, J. M.; Kirkegaard, R. H.; Michaelsen, T. Y.; Andersen, M. H.; Kondrotaite, Z.; Karst, S. M.; Dueholm, M. S.; Nielsen, P. H.; Albertsen, M. Connecting Structure to Function with the Recovery of over 1000 High-Quality

Metagenome-Assembled Genomes from Activated Sludge Using Long-Read Sequencing. *Nat. Commun.* **2021**, *12* (1), 2009. https://doi.org/10.1038/s41467-021-22203-2.
